# Chromatin marks govern mutation landscape of cancer at early stage of progression

**DOI:** 10.1101/074724

**Authors:** Kyungsik Ha, Hong-Gee Kim, Hwajin Lee

## Abstract

Accumulation of somatic mutations over time leads to tissue abnormalities, such as cancer. Somatic mutation rates vary across the genome in a cell-type specific manner, depending on the types of mutation processes^1–7^. Although recent studies have identified several determinants relevant to the establishment of the cancer mutation landscape^8–13^, these studies have yet to propose the major time point at which these factors come into play during cancer progression. Here, we analyzed whole genome sequencing data from two different types of precancerous tissues, monoclonal B-cell lymphocytosis and Barrett’s esophagus, and their matching cancer types along with 423 epigenetic features from normal tissues to determine the critical time point when chromatin features contribute to the formation of the somatic mutation landscape. Our analyses revealed that a subset of cell-of-origin associated chromatin features can explain more than 80% of the regional mutation variance for both types of precancerous tissues, comparable to the variance explained level for the genomes of matching cancer types. In particular, major significant chromatin features explaining the mutation landscape of Barrett’s esophagus and esophageal adenocarcinoma were derived from stomach tissues, indicating that mutation landscape establishment occurs mostly after environment-mediated epigenetic changes during gastric metaplasia. Analyses of the genome of esophageal squamous cell carcinoma tissues demonstrated that the proposed time point for mutation landscape establishment of Barrett’s esophagus and esophageal adenocarcinoma were specific to the occurrence of cell-type shift. Thus, our data suggest that the major time point for the mutation landscape establishment dictated by chromatin features is early in the process of cancer progression, and epigenetic changes due to environmental conditions at early stages can dramatically impact the somatic mutation landscape of cancer.

---

Recent advances in cancer genomics have so far revealed numerous somatic mutation landscapes for various cancer types, leading to a number of key findings. Identification of new driver gene mutations, deciphering clonal evolution structure and profiling tumor heterogeneity within and among different patients through examination of mutations, mainly at the gene level^1–7^, have successfully addressed the genes contributing to cancer progression and identified novel therapeutic targets. Beyond these gene-focused approaches, systematic analyses of mechanisms that could explain genomic regional variations in mutation rates across various cancer types could significantly extend our understanding about common contributors to the establishment of mutation landscapes before and during cancer progression. To this end, a number of studies have examined relationships between regional mutation frequencies across the genome and several types of features, including gene expression level, DNA sequence context, mutation profiles of nucleotide excision and mismatch repair genes, histone post-translational modifications, and open chromatin marks such as DNase-seq profiles^8–15^. Although these factors display high correlation with regional mutation rates, somatic mutation profiles used for the studies were limited to fully progressed tumors. Thus, it remains unknown whether the correlations between regional mutation frequencies and cell-of-origin chromatin marks are established either gradually during cancer progression or during a specific critical time period, either pre-or post-malignancy. Analyzing the mutation landscapes of precancerous, non-neoplastic tissues alongside matching cancer tissues could help to determine the major time points where chromatin marks shape the mutation landscape.

Here, we analyzed a total of 38 precancerous lesions including monoclonal B cell lymphocytosis (MBL) and Barrett’s esophagus (BE) (methods). Representative matching cancer types were also analyzed, corresponding to a total of 144 tumor samples from chronic lymphocytic leukemia (CLL) and esophageal adenocarcinoma (EAC). In addition, a total of 14 esophageal squamous cell carcinoma (ESCC) samples were analyzed to represent cancer without any defined precancerous stages during progression with a matching cell-of-origin.

We performed principal coordinate analysis (PCOA) to test whether the average mutation rate differences reported previously^16,17^ were reflected in the level of 1 megabase window regional mutation frequencies. Consistent with the differences in average mutation frequency, both MBL samples and CLL samples were indistinguishably located and formed separate clusters based on immunoglobulin heavy chain variable region (IGHV) mutation status, a key marker for distinguishing either naive-B cells or memory B cell origin for both MBL and CLL^16,18^. These results indicate that cell-of-origin differences might dictate the differences in regional mutation frequencies, rather than cancer progression-based status alone (Fig. 1a). In contrast, individual BE tissues formed clusters with the EAC tissues separate from the ESCC tissues, suggesting that the matching of cancer progression history might serve as a stronger factor than the cell-of-origin context itself (Fig. 1b). Collectively, these results show similarity in regional variation in mutation frequencies of precancerous tissues and matching cancer types, indicating that the effect of cell-of-origin context might be cancer-type dependent.

**Figure 1.**
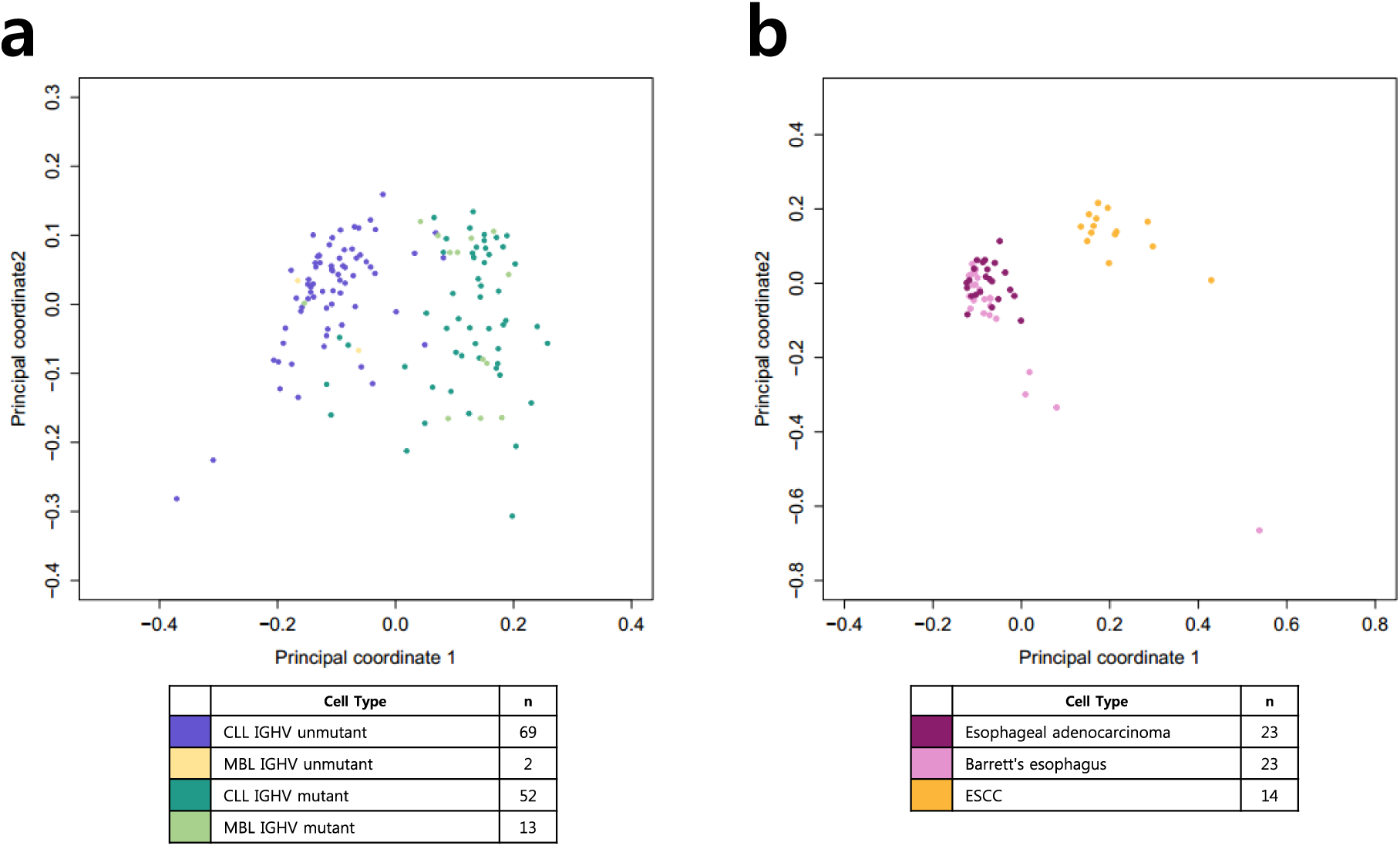
Principal coordinate analysis (PCOA) of individual cancer samples. (a) MBL and CLL with different IGHV mutation status. (b) Barrett’s esophagus, esophageal adenocarcinoma, and ESCC.

Whole-genome analyses of distinct cancer types depict cell-of-origin chromatin marks as the strongest feature explaining the cancer mutation landscape, with a number of proposed mechanisms^10^. Based on the IGHV mutation status-based clustering of MBL and CLL tissues in PCOA, we hypothesized that differential IGHV mutation status would correlate with distinct chromatin features explaining the regional mutation variation, and similar chromatin features would come up as significant when comparing IGHV mutation type-matching MBL and CLL genomes. To confirm the former part of the hypothesis, we first employed a random forest regression-based chromatin feature selection algorithm to identify significant chromatin features explaining the variance in regional mutation rates for different sample groups. Indeed, significant chromatin features explaining regional mutation variations were different between IGHV unmutant and groups (Extended Data Fig. 1a). Top-ranked chromatin features for both groups were derived from CD19-positive cells, which is expected since the CD19 marker cannot distinguish between naive and memory B cells. To further examine whether the differences in chromatin features were cell-type dependent, we performed chromatin feature selection after removing the 1Mbp regions containing IGHV mutation status-associated differential DNA methylation SNPs, which also highly overlaps with differential DNA methylation SNPs between naive and memory B cells^19–21^. This approach resulted in 3 out of 4 top significant chromatin features between the IGHV-mutant and unmutant groups (Extended Data Fig. 1b), implying that the differential chromatin features explaining mutation frequency landscapes of distinct IGHV mutation status might actually correlate with differences in cell-of-origin context. Next, we compared chromatin features that might explain regional mutation variations across the genomes of IGHV-mutation-status-matched MBL and CLL tissues. Due to the limits of sample size and average mutation rate of the samples, only IGHV-mutant MBL and CLL genomes were subjected to further analyses. Notably, the top two ranked chromatin features were identical between IGHV-mutant MBL and CLL samples (Fig. 2a), implicating that the subset of chromatin marks might commonly dictate the formation of regional mutation landscape for both pre-cancerous tissues and matching cancer type. Additional examination of simple correlation between regional mutation frequency and histone modification levels derived from CD19-positive cells at the 1 megabase-level revealed marginal differences between MBL and CLL tissues (Fig. 2b and Extended Data Fig. 2a). The correlation between the CD19 DNase1-seq profile and regional mutation frequency was higher for CLL than MBL for chromosome 2 (Fig. 2c) and other chromosomes (Extended Data Fig. 3a), but this finding might be due to the different number of samples between MBL and CLL, as the correlation score for MBL at chromosome 2 was highly similar to the correlation scores for CLL (0.76 vs. 0.75) after sample-number matching. These results demonstrate that the cell-of-origin chromatin context, defined by the IGHV mutation status, serves a major role in shaping the mutation landscape of both MBL and CLL tissues, suggesting that the cell-of-origin chromatin landscape could govern the establishment of the somatic mutation landscape for CLL early in cancer progression, even before the precancerous cell type, MBL, is apparent.

**Figure 2.**
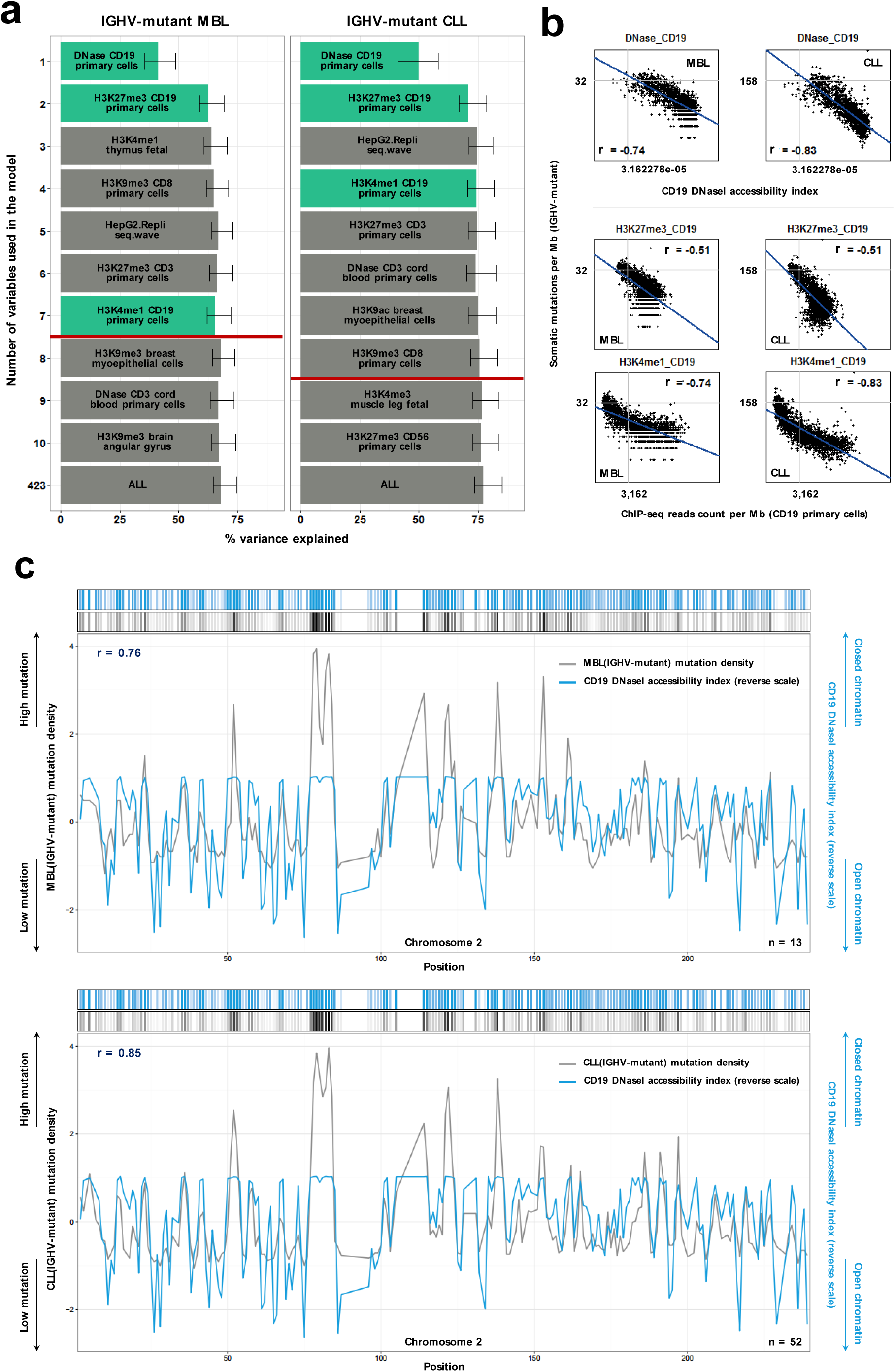
Cell-of origin chromatin features representing association with the regional mutation frequency of CLL and the corresponding precancerous cell type, MBL. (a) Random forest regression-based chromatin feature selection in relations to the regional mutation frequency of IGHV-mutant MBL and CLL samples. Each chromatin feature is ranked by importance value, and variance explained scores are represented by bar length. Error bars demonstrate minimum and maximum values derived from 1,000 repeated simulations. Red lines display variance explained scores determined by 423 features - 1 SEM, and CD19 chromatin features are green-colored. (b) Univariate correlation between CD19 chromatin features that displayed significance in the feature selection models and the regional mutation density of IGHV-mutant MBL or CLL. Spearman’s rank correlations (*r*) are shown on each plot. (c) The density plot for regional mutation density of IGHV-mutant MBL or CLL and CD19 DNase1 accessibility index (reverse scale) across the full chromosome 2. Spearman’s rank correlations (γ) are shown on each plot.

Cell type shift, represented as gastric metaplasia, is one of the main hallmarks in the development of BE^22^. Thus, one could assume that the critical time point for the establishment of the mutation landscape for BE could be either before or during the course of cell type shift, or after its completion. Chromatin feature selection analysis of the mutation landscape of BE and EAC tissues confirmed that high-ranked chromatin features were derived from the stomach tissue type (Extended Data Fig. 4) for both tissues, without any significant esophageal chromatin features. Simple correlation between regional mutation frequency and histone modification marks from stomach and esophagus tissues revealed marginal differences between BE and EAC tissues (Extended Data Fig. 2b, c), and this pattern was also consistent with the correlation to stomach tissue DNase I hypersensitivity profile (Extended Data Fig. 3b). Moreover, six features covering all stomach chromatin features subjected to the feature selection analysis solely explained over 80% of the regional mutation variance for both BE and EAC tissues, which is unlikely to be random (p value < 2.2e-16) (Extended Data Fig. 5). These results imply that the major time point of mutation landscape establishment for BE is most likely to be after the cell type shift into stomach mucosa-like cells. Chromatin feature selections on subgroups of somatic mutations for BE and EAC based on overlap and uniqueness of the mutations shared common top-ranked stomach chromatin features (Fig. 3a). In addition, chromatin feature selection on sample subgroups with respect to dysplasia grade revealed that the top features all originated from stomach tissue (Extended Data Fig. 6) and the variance explained level for all of the dysplasia-based subgroups using six stomach tissue chromatin features were similar to the variance explained level using all 423 chromatin features (Fig. 3a). This finding was consistent with the high correlation to stomach tissue DNase I hypersensitivity profile (Extended Data Fig. 3c). From all of these results, we infer an early time point for establishment of the mutation landscape for EAC, even prior to the occurrence of dysplasia for BE, but most likely after epigenetic changes due to gastric metaplasia.

**Figure 3.**
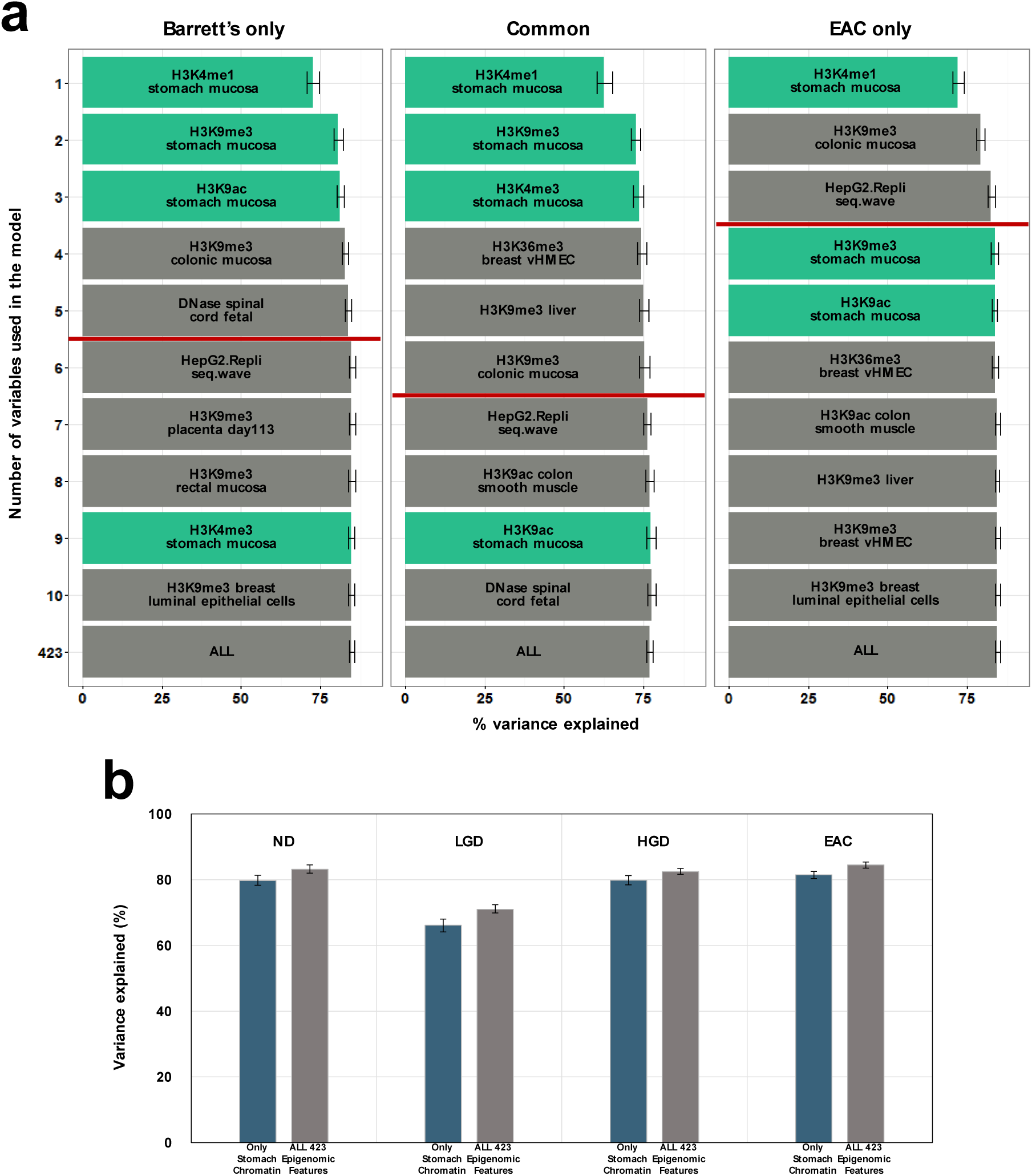
Regional mutation frequency landscape of Barrett’s esophagus and matching esophageal adenocarcinoma are affected by cell-type-shift-associated epigenetic changes. (a) Chromatin feature selection based on the commonality of mutations in paired samples of Barrett’s esophagus and esophageal adenocarcinoma. Barrett’s only: mutations observed only in the Barrett’s esophagus genome, Common: mutations observed in common for both Barrett’s esophagus and esophageal adenocarcinoma genomes, EAC only: mutations observed solely in the esophageal adenocarcinoma genome. (b) Bar graph representing average variance explained scores using either stomach chromatin features (navy) or all 423 epigenomic features (gray). ND: no dysplasia, LGD: low-grade dysplasia, HGD: high-grade dysplasia, EAC: esophageal adenocarcinoma. Error bars demonstrate minimum and maximum values derived from 1,000 repeated simulations.

To ensure that the chromatin features shaping the mutation landscape of BE and EAC were not common to any esophageal cancer type, we analyzed the genome of ESCC, another cancer type derived from the esophageal squamous epithelium without any precancerous stages with cell type shift. Although the regional mutation frequency of ESCC correlated with histone modification marks from stomach and esophagus tissues in a similar manner (Extended Data Fig. 2d), chromatin feature selection revealed a subset of squamous cell type and esophagus chromatin features that were significant and distinct from BE and EAC (Extended Data Fig. 7). Moreover, measuring the level of variance explained values per tissue or cell type categories showed stomach chromatin features to be the strongest ones for BE and EAC, reaching higher than 90% of the variance level explained by the 423 total chromatin features, whereas esophageal chromatin features were dominant for ESCC (Fig. 4). Notably, the variance explained values for each category displayed non-significant relationship with simple correlations between the chromatin marks from different tissue or cell types (BE *r_s_* = 0.36, EAC *r_s_* = 0.36, ESCC *r_s_* = 0.18). These results imply a distinct process of mutation landscape establishment for these cancer types that varies depending on the presence of precancerous tissues with cell-type shifts.

**Figure 4.**
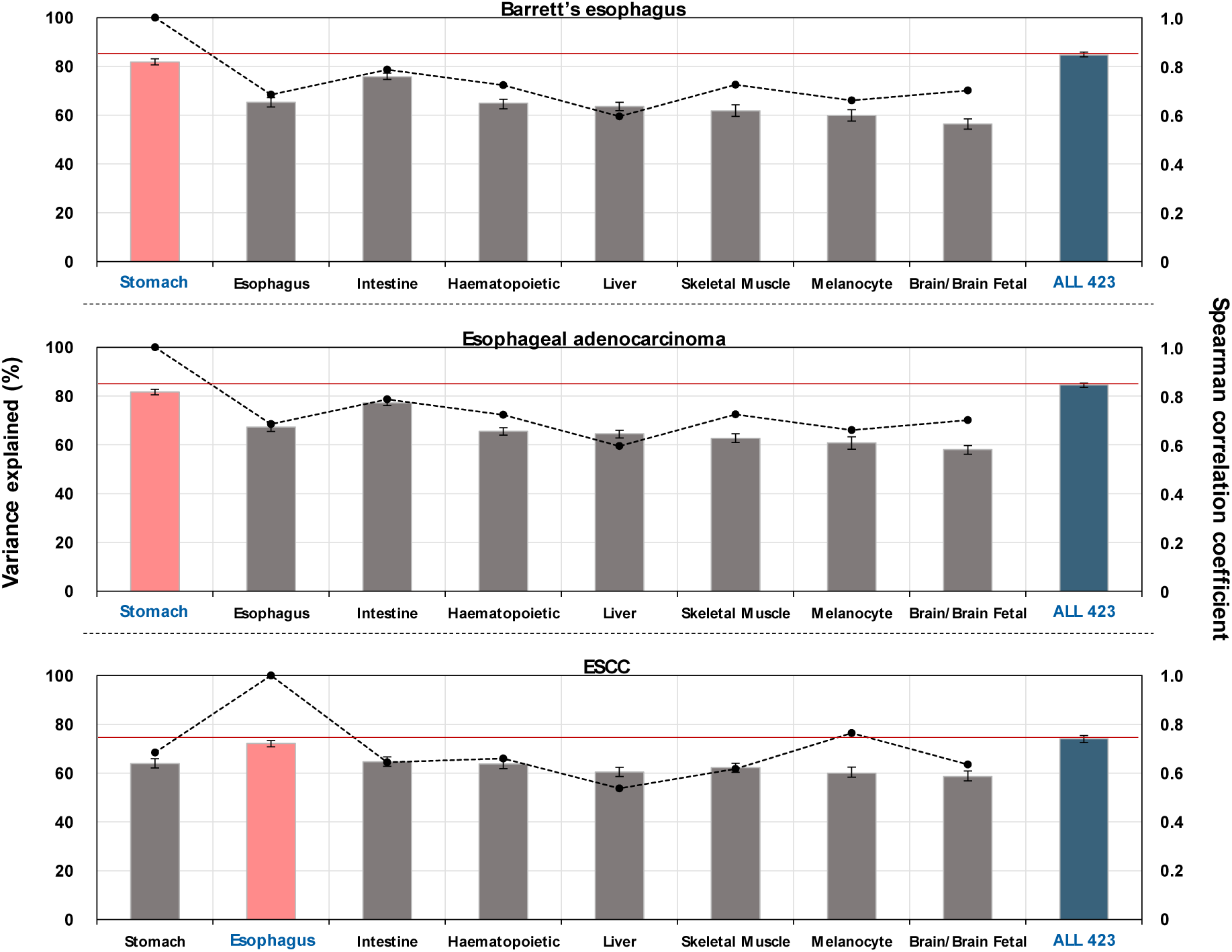
Regional mutation frequency landscape of esophageal squamous cell carcinoma demonstrates the uniqueness of significant chromatin features associated with the Barrett’s esophagus and esophageal adenocarcinoma genomes. Average variance explained scores for pre-cancerous or matching cancer genomes were separately calculated using the tissue or cell type-based subgroup-classified chromatin features. The pink panel represents subgroups with the highest variance explained score for each cell type. The red line indicates the variance explained score when using all 423 epigenomic features. Dots represent the Spearman’s rank correlations (r) of chromatin features between the highest variance explained-scored subgroup and the remaining subgroups. Error bars demonstrate minimum and maximum values derived from 1,000 repeated simulations.

In conclusion, our data suggest that the major time point for the establishment of the mutation landscape governed by chromatin marks could be early, even prior to the phenotypic emergence of precancerous tissues. Results from BE and EAC also raise the possibility that epigenetic changes due to environmental insults, represented as a cell type shift, could serve as a primary role by affecting the course of the establishment of the mutation landscape (Extended Data Fig. 8). Further comprehensive studies to decipher the mutation landscape of other precancerous tissues with metaplasia and discover the exact mechanisms controlling the timing of mutation landscape establishment would lead to a better understanding of the effect of epigenetic marks on shaping the precancerous tissues and matching cancer genome and help identify possible biomarkers for early-stage detection of cancer.

## METHODS

### Data

For the purposes of our project, we used somatic mutation data from CLL, MBL, BE, EAC, and ESCC tissues. In the case of CLL and MBL genome data, total mutations were acquired from Supplementary Table 2 of the publication^16^, consisting of 136 samples (13 IGHV-mutant MBL, 2 IGHV-unmutant MBL, 52 IGHV-mutant CLL and 69 IGHV-unmutant CLL). In the case of Barrett’s esophagus, esophageal adenocarcinoma, and ESCC, data use were authorized from ICGC (http://icgc.org) and BGI (http://www.genomics.cn/) before use. A total of 23 pairs of Barrett’s esophagus and matching esophageal adenocarcinoma genomics data^17^ were authorized from ICGC and genome data of 14 ESCC samples^23^ were acquired from BGI. These data sets were subsequently analyzed following the standard GATK pipeline (https://www.broadinstitute.org/gatk/) and somatic variants were called using the MuTect algorithm^24^ (https://www.broadinstitute.org/cancer/cga/mutect).

Epigenomics and chromatin data were from the NIH Roadmap Epigenomics Mapping Consortium^25^ and ENCODE^26^. NIH Roadmap Epigenomics data were accessible from the NCBI GEO series GSE18927, referring to the University of Washington Human Reference Epigenome Mapping Project.

To calculate the regional mutation density and mean signal of chromatin features, all autosomes were split in 1-Mbp regions followed by filtering out regions containing centromeres, telomeres and low quality unique mappable base pairs. To determine regional mutation density and histone modification profiles, we counted the total number of somatic mutations or ChIP-seq reads per each 1 megabase region. For analyzing the DNase I hypersensitivity and Repli-seq data, scores of DNase I peaks and replication were calculated per each 1 megabase region. For somatic mutations, ChIP-seq data and DNase I hypersensitivity data, BEDOPS^27^ was employed to calculate the frequency and scores per each 1Mbp region.

### Principal coordinate analysis

Principal coordinate analysis was used to represent differences in mutation frequency distribution among the individual samples. A dissimilarity matrix was built using 1 – Pearson correlation coefficient across all samples. Each sample location was assigned in a two-dimensional space using this matrix.

### Feature selection based on random forest regression

A random forest regression-based feature selection algorithm was performed as described^10^ with modifications. Briefly, the training set for each tree was constructed, followed by using out-of-bag data to estimate the mean squared error. Thus, there was no need to perform additional tests for error evaluation. Out-of-bag data were also used to estimate the importance of each variable. In each out-of-bag case, the values corresponding to each variable were randomly permuted, then tested to each tree. Subtracting the score of the mean squared error between the untouched out-of-bag data cases and the variable-m-permuted cases, the raw importance score of variable m was measured. By calculating the average score of variable m in the entire tree, the rank of importance for each variable was determined. A total of 1,000 random forest trees were employed to predict mutation density using a total of 423 chromatin features. Every random forest model was repeated 1,000 times.

After the random forest algorithm step, greedy backward elimination was performed to select the top 20 chromatin variables. Subsequent removal of the lowest rank variable was done to calculate the variance explained value measurements for each variable. To conduct feature selection on all of the samples corresponding to the particular pre-cancerous tissues or cancer types, mutation density was calculated by adding samples in each case. However, a number of particular analyses employed the subgrouping of samples. In the case of chromatin feature selection assessing the effects of differential DNA methylation between IGHV-mutants and unmutants (Extended Data Fig. 1b), a total of 935 regions containing differentially methylated CpGs^20^ were removed prior to the analysis. To perform feature selection classified by differential dysplasia states (Extended Data Fig. 6), samples were divided into 3 groups: 17 samples of no dysplasia, 3 samples of low-grade dysplasia and 2 samples of high-grade dysplasia. In the case of feature selection after subgrouping for distinct and common mutations (Fig. 3a), all mutations in paired-samples of BE and EAC were divided into 3 different groups: Barrett’s only, EAC only, and common mutations.

### Analysis of mutation frequency variance explained by chromatin features

To examine the effect of a particular cell-type specific chromatin context on explaining regional variability of mutation density across the genome, chromatin features were subgrouped based on the feature selection algorithm. To study the differences in variance explained values among distinct cell types, 8 groups were categorized (Fig. 4). Each group included 5 chromatin markers common among the 8 cell-type based groups: H3K27me3, H3K36me3, H3K4me1, H3K4me3 and H3K9me3. Random selection of 6 chromatin features were either from all of the 423 features or 417 features (excluding stomach mucosa chromatin features) (Extended Data Fig. 5). Random selection of chromatin features was repeated 1,000 times, then the average variance explained values and permutation distributions were obtained.

## Acknowledgements

This work was supported by Basic Science Research Program through the National Research Foundation of Korea(NRF) funded by the Ministry of Science, ICT and future Planning (NRF-2014R1A2A1A11049728). The manuscript was proofread by the Dental Research Institute of Seoul National University. We thank Dr. Paz Polak for providing sample epigenomics data with helpful comments. We also like to thank Minji Kim and Yongju Lee for the involvement of designing and fine-tuning the individual figures.

## Author Contribution

H.L. provided the original idea. H.L., K.H. and H.K. led the overall project. K.H and H.L analyzed the data and contributed to scientific discussions. H.L., K.H. and H.K wrote the manuscript.

## Author Information

H.L is currently working at Celltrion, Inc. to fulfill his military service duty, but conducted the current research without any conflict of financial interests. Other authors declare no competing financial interests. Readers are welcome to comment on the online version of the paper. Correspondence and requests for materials should be addressed to H.K. or H.L.

---

**Extended Data Figure 1.**
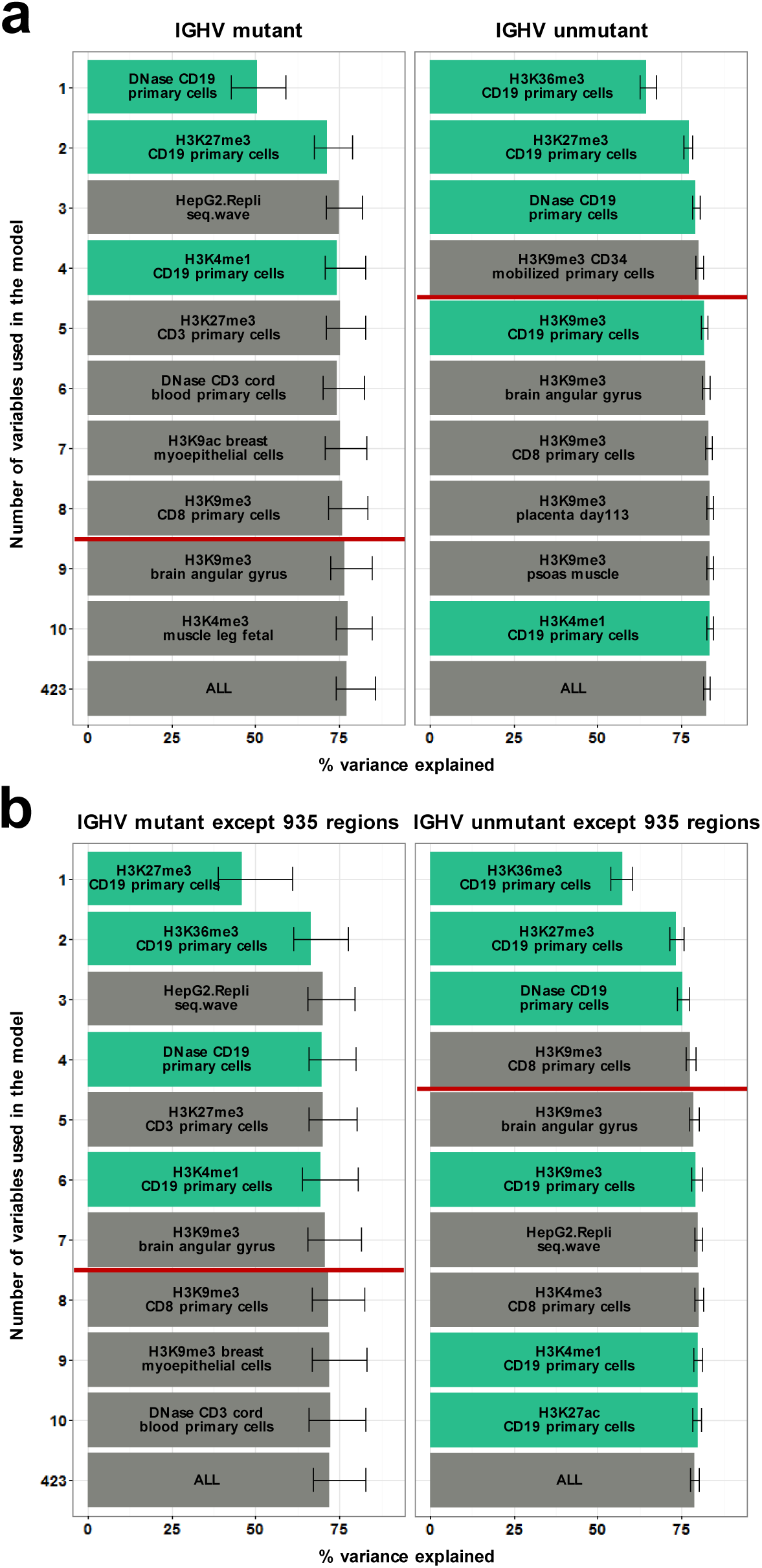
Random forest regression-based chromatin feature selection using IGHV-mutant sample groups or IGHV-unmutant sample groups. (a) IGHV-mutant vs. IGHV-unmutant samples with all 1 megabase genomic regions. (b) IGHV-mutant vs. IGHV-unmutant samples without 935 of the 1 megabase genomic regions corresponding to regions containing differentially methylated CpGs between IGHV-mutant and IGHV-unmutant CLL samples.

**Extended Data Figure 2.**
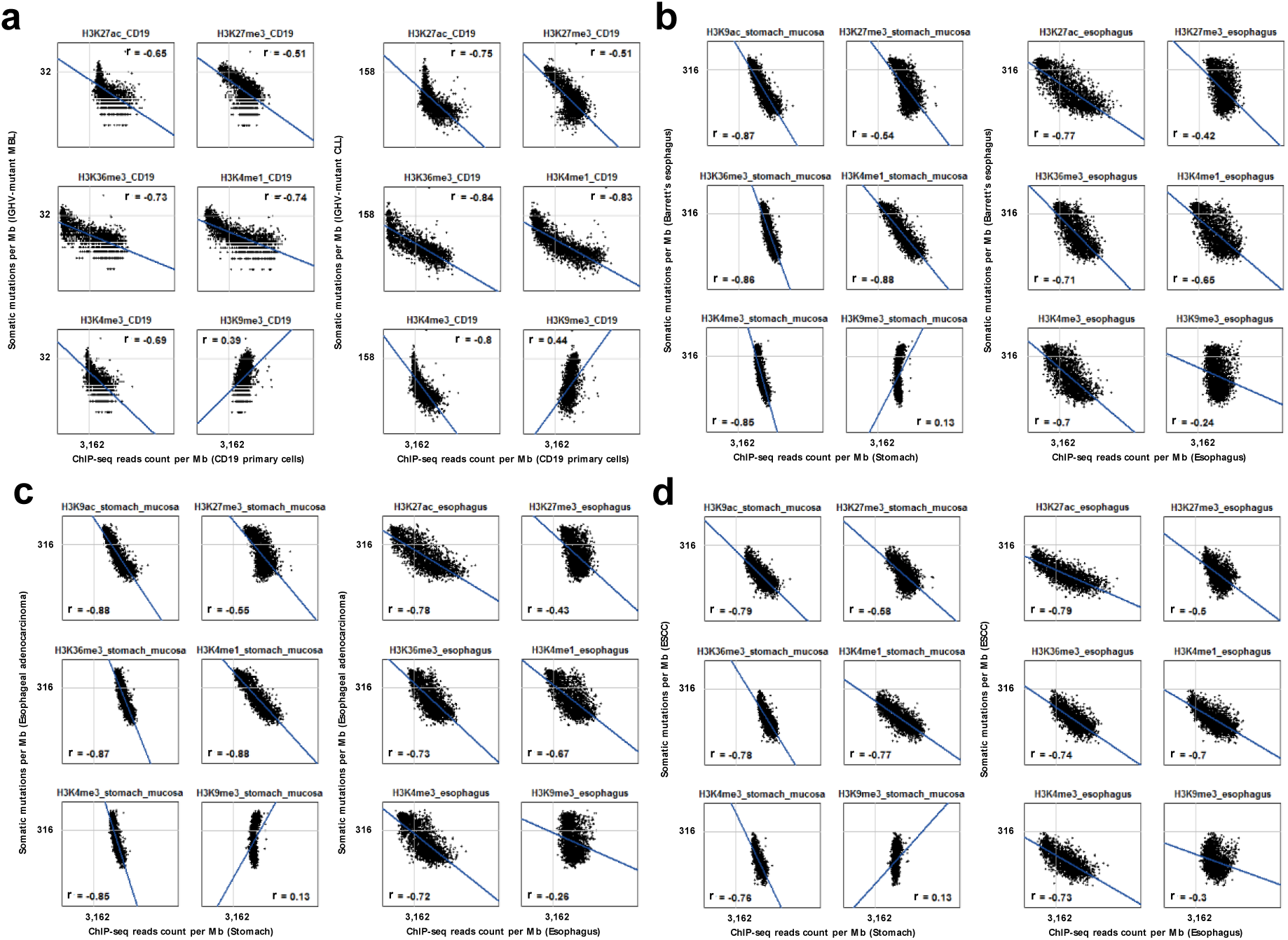
Correlation plots between regional mutation density and cell-type matching chromatin features. (a) Mutation density of IGHV-mutant MBL or CLL versus CD19 chromatin features. (b) Mutation density of Barrett’s esophagus versus stomach mucosa or esophagus chromatin features. (c) Mutation density of esophageal adenocarcinoma versus stomach mucosa or esophagus chromatin features. (d) Mutation density of ESCC versus stomach mucosa or esophagus chromatin features.

**Extended Data Figure 3.**
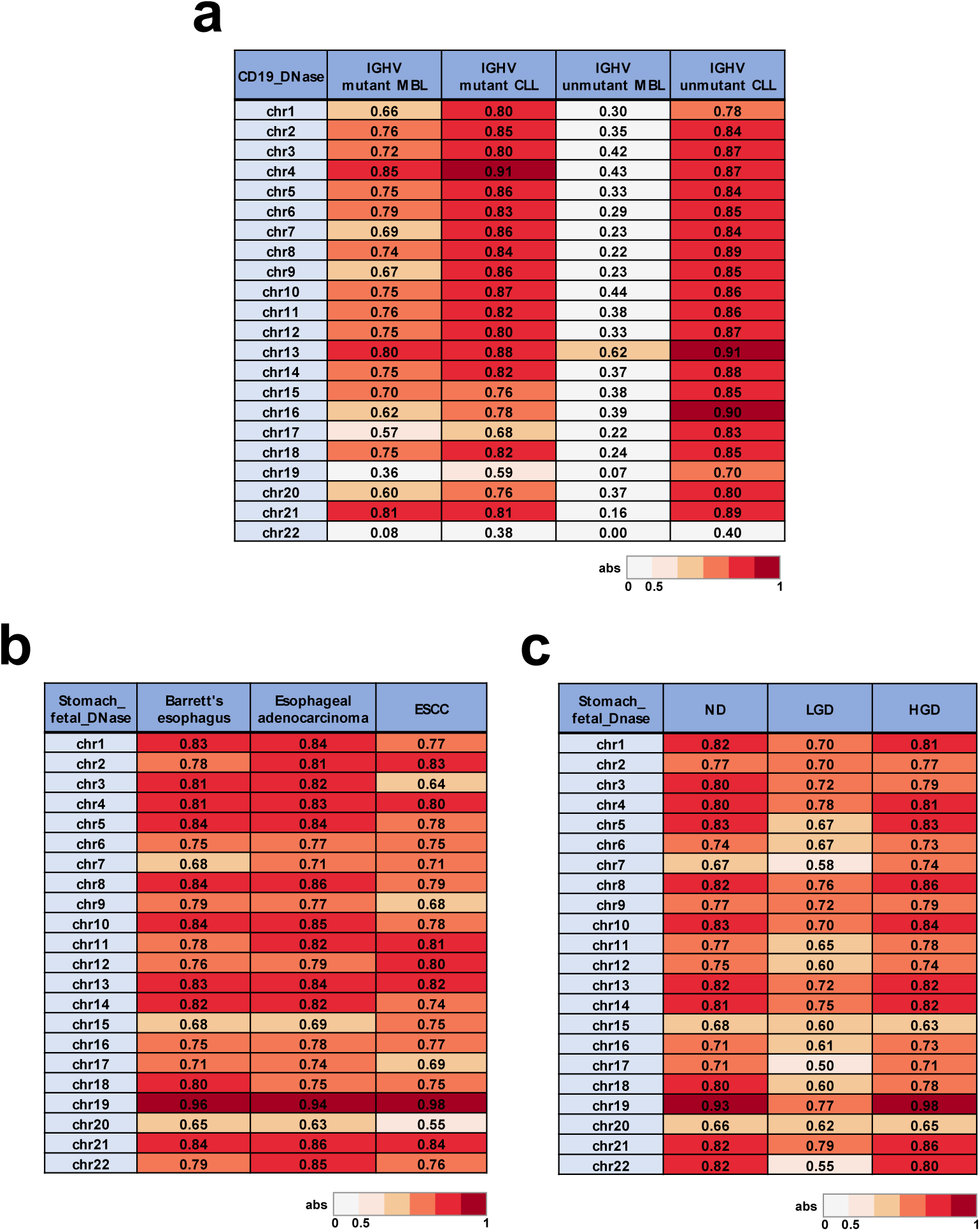
Spearman’s rank correlation (γ) between regional mutation density and chromatin accessibility index across the different chromosomes. (a) MBL and CLL with different IGHV mutation status. (b) Barrett’s esophagus, esophageal adenocarcinoma and ESCC. (c) Subgroups of Barrett’s esophagus classified by dysplasia states.

**Extended Data Figure 4.**
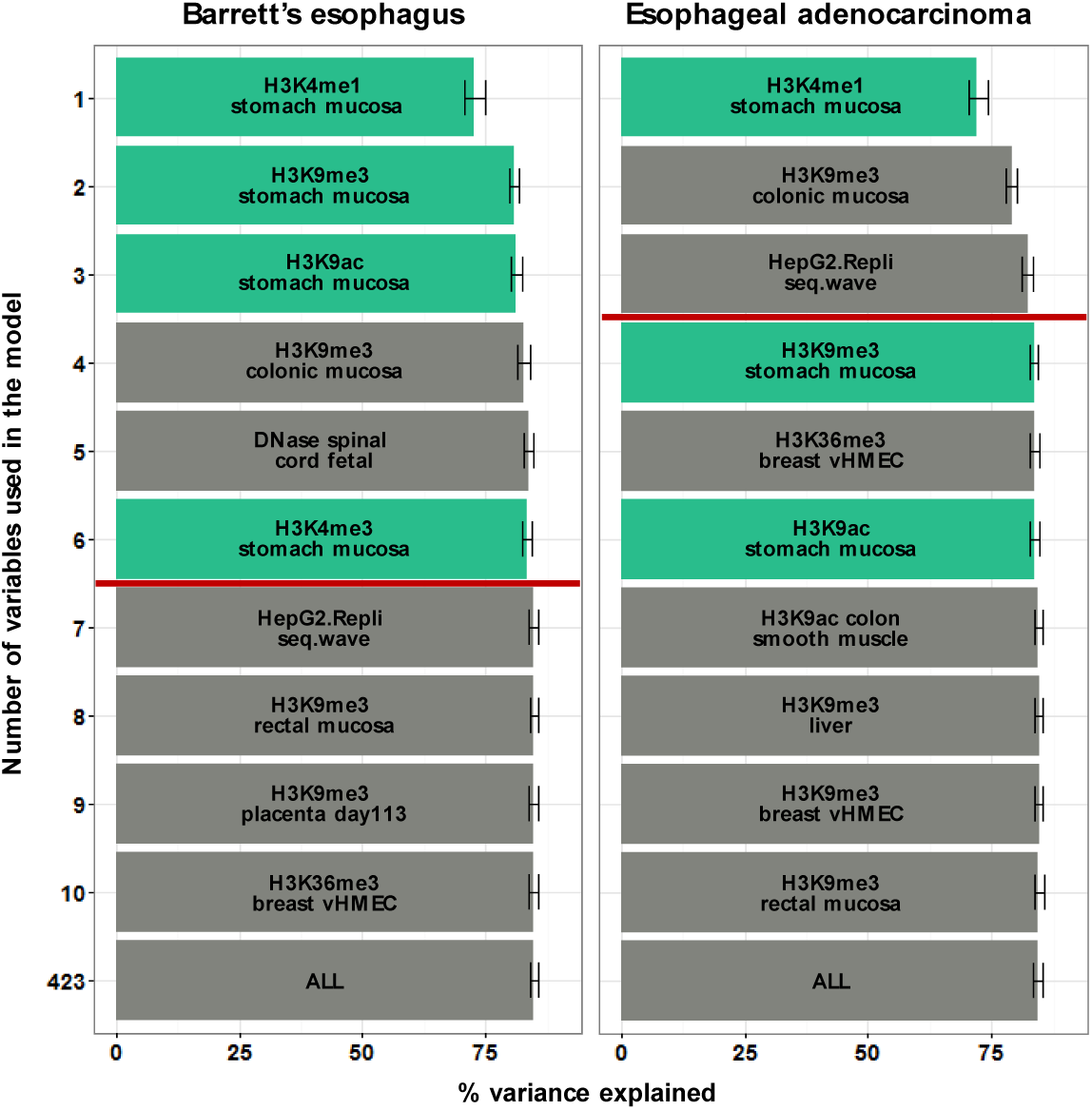
Chromatin feature selection in relation to the regional mutation frequency of Barrett’s esophagus and esophageal adenocarcinoma. Chromatin features of the stomach mucosa are green-colored.

**Extended Data Figure 5.**
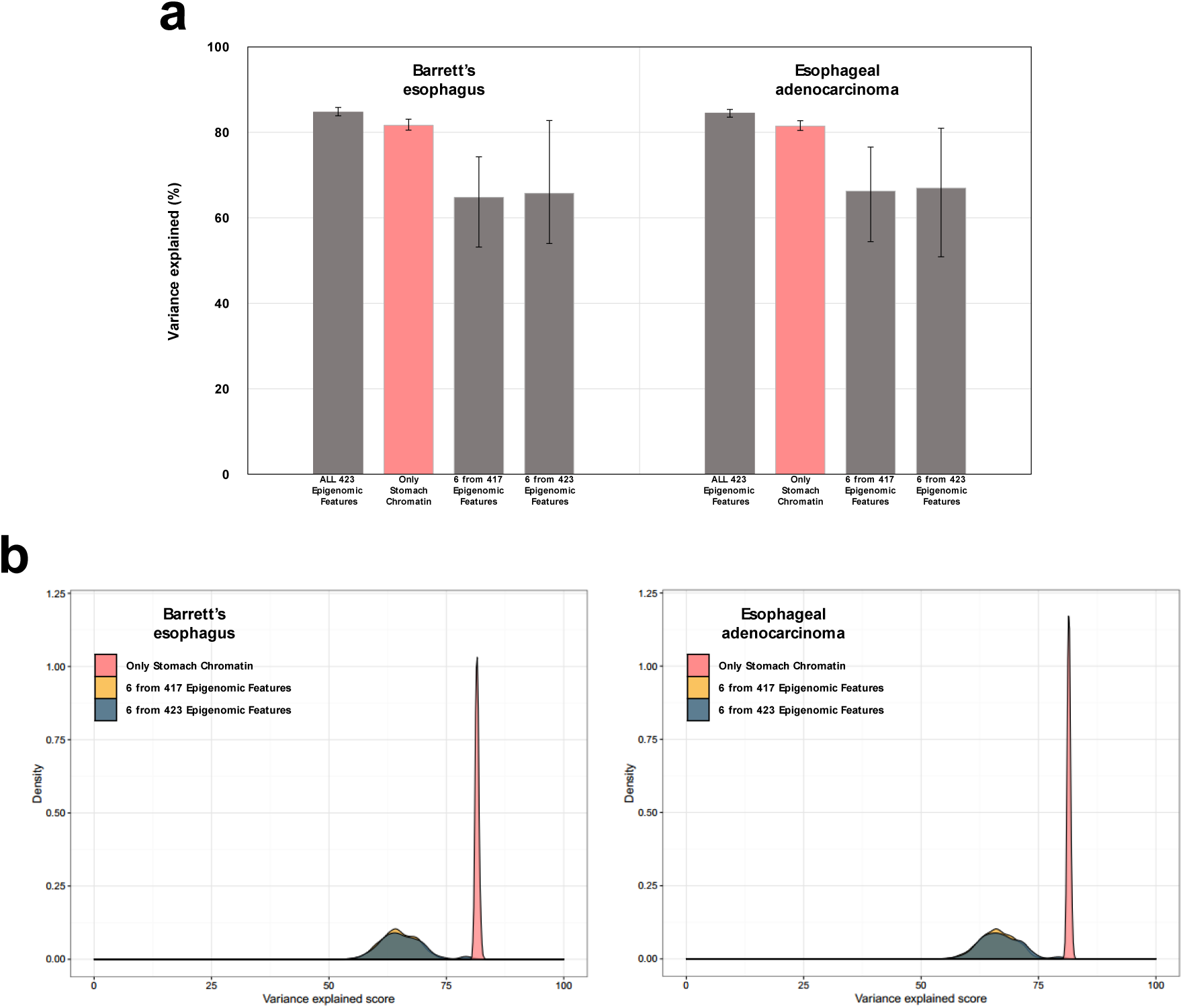
Comparison of variance explained scores using either stomach chromatin features or groups of randomly selected chromatin features. Stomach chromatin group represents a total of 6 chromatin features from stomach tissue. A total of 417 and 423 chromatin groups displayed 6 randomly selected chromatin features from either 417 or 423 features. The difference between 417 and 423 features was the presence or absence of stomach chromatin features. (a) Average variance explained scores using 3 different chromatin groups or all of the 423 features. Error bars demonstrate minimum and maximum values derived from 1,000 repeated simulations. (b) Distribution of variance explained scores for the group of 6 randomly selected chromatin features from either 417 or 423 chromatin features with 1,000 permutations. Pink-colored distributions represent average variance explained score of stomach chromatin features.

**Extended Data Figure 6.**
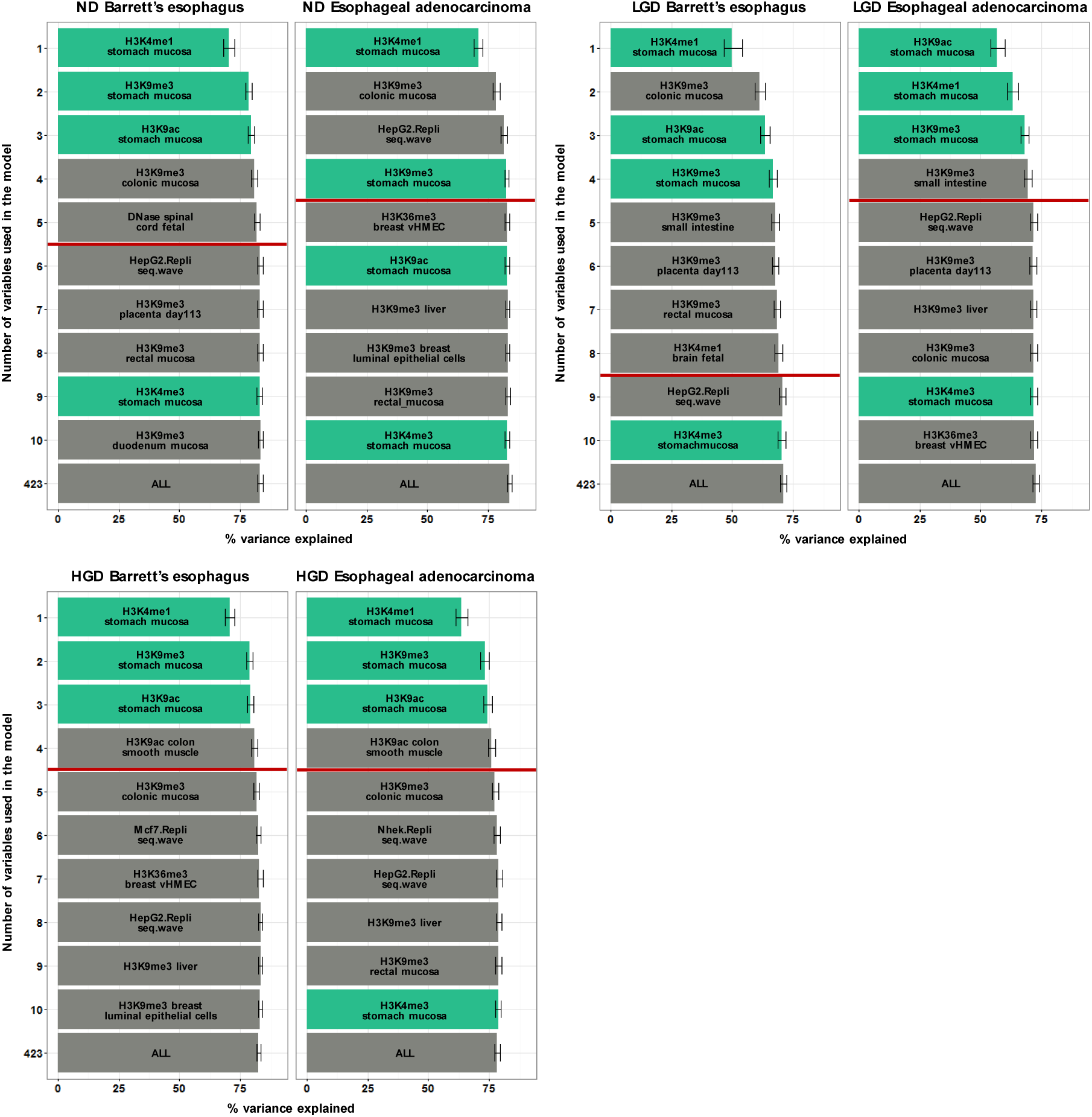
Feature Selection in Barrett’s esophagus and esophageal adenocarcinoma classified by dysplasia status.

**Extended Data Figure 7.**
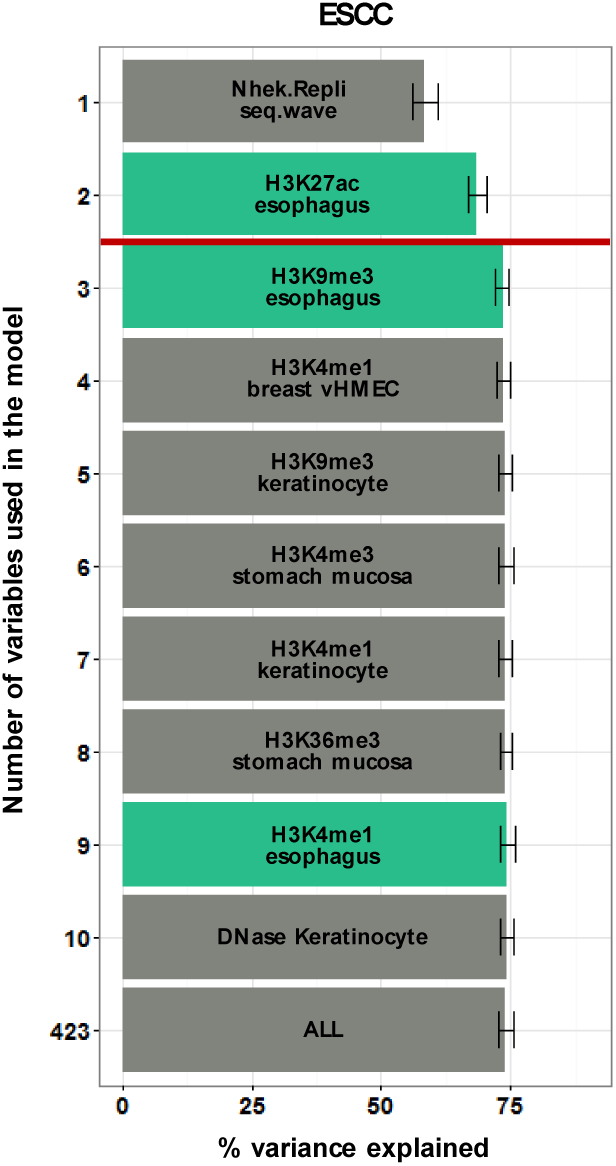
Chromatin feature selection in relation to the regional mutation frequency of ESCC samples. Chromatin features of the esophagus are green-colored.

**Extended Data Figure 8.**
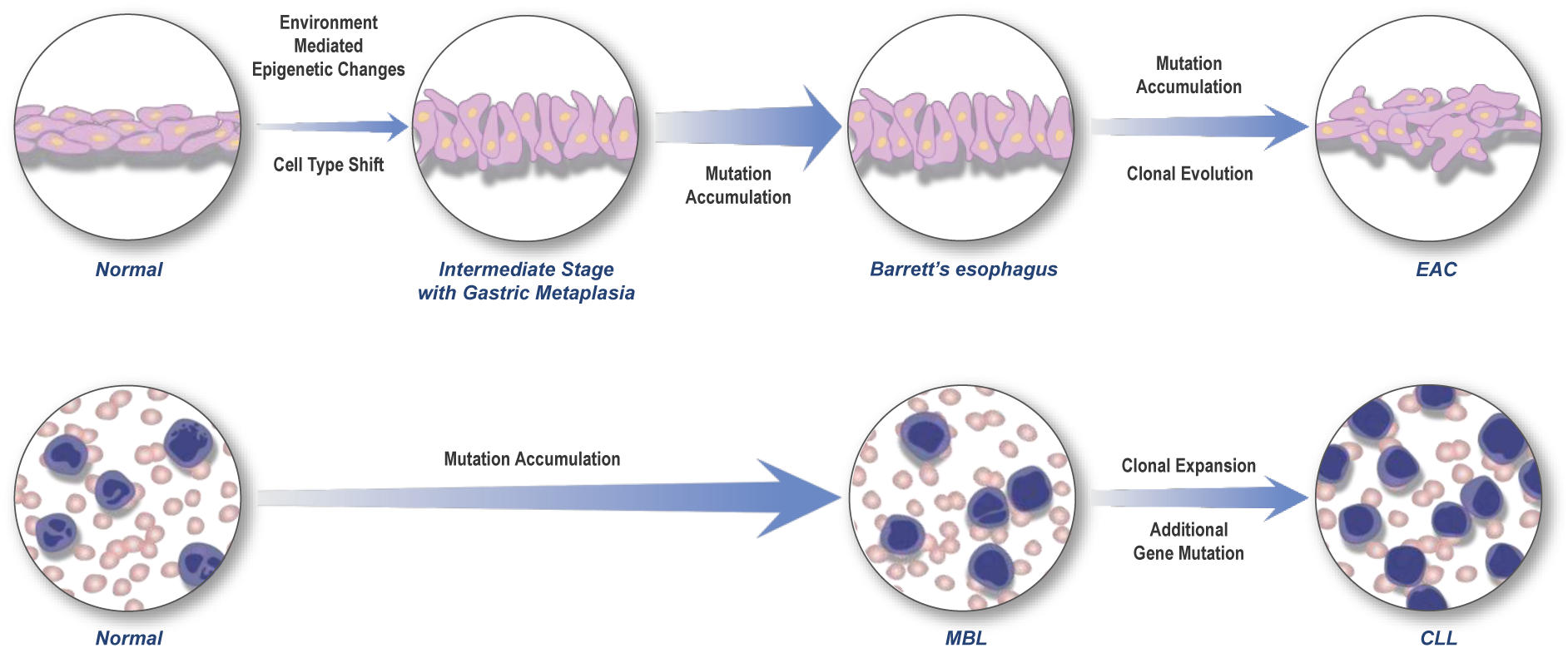
Proposed model showing the major time point for the establishment of the mutation landscape with respect to chromatin features.

## References

1. Alexandrov, L. B. et al. Signatures of mutational processes in human cancer. Nature 500, 415–421, doi:10.1038/nature12477 (2013).

2. Hodgkinson, A., Chen, Y. & Eyre-Walker, A. The large-scale distribution of somatic mutations in cancer genomes. Hum Mutat 33, 136–143, doi:10.1002/humu.21616 (2012).

3. Kan, Z. et al. Diverse somatic mutation patterns and pathway alterations in human cancers. Nature 466, 869–873, doi:10.1038/nature09208 (2010).

4. Kandoth, C. et al. Mutational landscape and significance across 12 major cancer types. Nature 502, 333–339, doi:10.1038/nature12634 (2013).

5. Lawrence, M. S. et al. Mutational heterogeneity in cancer and the search for new cancer-associated genes. Nature 499, 214–218, doi:10.1038/nature12213 (2013).

6. Martincorena, I. & Campbell, P. J. Somatic mutation in cancer and normal cells. Science 349, 1483–1489, doi:10.1126/science.aab4082 (2015).

7. Schaefer, M. H. & Serrano, L. Cell type-specific properties and environment shape tissue specificity of cancer genes. Sci Rep 6, 20707, doi:10.1038/srep20707 (2016).

8. Liu, L., De, S. & Michor, F. DNA replication timing and higher-order nuclear organization determine single-nucleotide substitution patterns in cancer genomes. Nat Commun 4, 1502, doi:10.1038/ncomms2502 (2013).

9. Polak, P. et al. Reduced local mutation density in regulatory DNA of cancer genomes is linked to DNA repair. Nat Biotechnol 32, 71–75, doi:10.1038/nbt.2778 (2014).

10. Polak, P. et al. Cell-of-origin chromatin organization shapes the mutational landscape of cancer. Nature 518, 360–364, doi:10.1038/nature14221 (2015).

11. Schuster-Bockler, B. & Lehner, B. Chromatin organization is a major influence on regional mutation rates in human cancer cells. Nature 488, 504–507, doi:10.1038/nature11273 (2012).

12. Stamatoyannopoulos, J. A. et al. Human mutation rate associated with DNA replication timing. Nat Genet 41, 393–395, doi:10.1038/ng.363 (2009).

13. Woo, Y. H. & Li, W. H. DNA replication timing and selection shape the landscape of nucleotide variation in cancer genomes. Nat Commun 3, 1004, doi:10.1038/ncomms1982 (2012).

14. Supek, F.&Lehner, B. Differential DNA mismatch repair underlies mutation rate variation across the human genome. Nature 521, 81–84, doi:10.1038/nature14173 (2015).

15. Thurman, R. E. et al. The accessible chromatin landscape of the human genome. Nature 489, 75–82, doi:10.1038/nature11232 (2012).

16. Puente, X. S. et al. Non-coding recurrent mutations in chronic lymphocytic leukaemia. Nature 526, 519–524, doi:10.1038/nature14666 (2015).

17. Ross-Innes, C. S. et al. Whole-genome sequencing provides new insights into the clonal architecture of Barrett's esophagus and esophageal adenocarcinoma. Nat Genet 47, 1038–1046, doi:10.1038/ng.3357 (2015).

18. Fabbri, G. & Dalla-Favera, R. The molecular pathogenesis of chronic lymphocytic leukaemia. Nat Rev Cancer 16, 145–162, doi:10.1038/nrc.2016.8 (2016).

19. Cahill, N. et al. 450K-array analysis of chronic lymphocytic leukemia cells reveals global DNA methylation to be relatively stable over time and similar in resting and proliferative compartments. Leukemia 27, 150–158, doi:10.1038/leu.2012.245 (2013).

20. Kulis, M. et al. Epigenomic analysis detects widespread gene-body DNA hypomethylation in chronic lymphocytic leukemia. Nat Genet 44, 1236–1242, doi:10.1038/ng.2443 (2012).

21. Oakes, C. C. et al. DNA methylation dynamics during B cell maturation underlie a continuum of disease phenotypes in chronic lymphocytic leukemia. Nat Genet 48, 253–264, doi:10.1038/ng.3488 (2016).

22. Hayakawa, Y., Sethi, N., Sepulveda, A. R., Bass, A. J. & Wang, T. C. Oesophageal adenocarcinoma and gastric cancer: should we mind the gap? Nat Rev Cancer 16, 305–318, doi:10.1038/nrc.2016.24 (2016).

23. Zhang, L. et al. Genomic analyses reveal mutational signatures and frequently altered genes in esophageal squamous cell carcinoma. Am J Hum Genet 96, 597–611, doi:10.1016/j.ajhg.2015.02.017 (2015).

24. Cibulskis, K. et al. Sensitive detection of somatic point mutations in impure and heterogeneous cancer samples. Nat Biotechnol 31, 213–219, doi:10.1038/nbt.2514 (2013).

25. Roadmap Epigenomics, C. et al. Integrative analysis of 111 reference human epigenomes. Nature 518, 317–330, doi:10.1038/nature14248 (2015).

26. Consortium, E. P. An integrated encyclopedia of DNA elements in the human genome. Nature 489, 57–74, doi:10.1038/nature11247 (2012).

27. Neph, S. et al. BEDOPS: high-performance genomic feature operations. Bioinformatics 28, 1919–1920, doi:10.1093/bioinformatics/bts277 (2012).

